# Inflammation drives age-induced loss of tissue resident macrophages

**DOI:** 10.1101/2022.10.02.510513

**Authors:** Kémy Adé, Javier Saenz Coronilla, Dorian Obino, Tobias Weinberger, Caroline Kaiser, Sébastien Mella, Cheng Chen, Lida Katsimpardi, Catherine Werts, Han Li, Pascal Dardenne, Yvan Lallemand, Elisa Gomez Perdiguero

## Abstract

Low-grade chronic systemic inflammation, or inflammageing, is a hallmark of ageing and a risk factor for both morbidity and mortality in elderly people. Resident macrophages are tissue homeostasis sentinels that are embedded in their tissue of residence since embryonic development, thus been exposed to cumulative tissue insults throughout life. Therefore, resident macrophages, among other immune cells, emerge as potential key contributors to age-associated tissue dysfunction. Contrary to what is currently postulated, we demonstrate here that the pool of embryo-derived resident macrophages exhibits an age-dependent depletion in liver, and other solid organs and that they are not replaced by Hematopoietic Stem Cell (HSCs)-derived monocytes throughout life. Further, we demonstrate that gradual, cumulative inflammation during ageing induces this specific loss of tissue resident macrophages. Preserving a “youthful” density of resident macrophages attenuates classical hallmarks of liver age-associated dysfunction.

**Summary:** The pool of embryo-derived resident macrophages dwindles with age in most tissues, without compensation from Hematopoietic Stem Cell (HSC)-derived cells. This loss is not due to impaired self-renewal in old tissues but rather to increased cell death, which is driven by sustained inflammation. Attenuating inflammation sensing during ageing prevents age-induce macrophage loss and improves hallmarks of liver ageing.

## Introduction

Ageing is associated with excessive innate immune cell production, defective adaptive immune responses, and systemic low grade chronic inflammation^1,2^. While the frequency of circulating innate immune cells is increased in aged animals, there are conflicting results regarding tissue-specific myeloid cells^3–5^. Among them, two lineages of macrophages coexist in most adult tissues: embryo-derived Hematopoietic Stem Cell (HSC)-independent “resident” macrophages and HSC-derived monocytes/macrophages^6^. Pathological situations have shed light on the impact of ontogeny on tissue macrophage functions^7–10^. However, the homeostatic maintenance and functions of resident macrophages are still poorly understood, in particular in the context of ageing. Whether HSC-derived macrophages can compensate for resident macrophage loss is a subject of intense investigation^11,12^. Given that the progressive replacement of heart resident macrophages with HSC-derived cells over time^13,14^ has been linked to transcriptomic changes that may augment age-associated cardiac fibrosis and deleterious outcomes following cardiac injury^15,16^, a shift in ontogeny among tissue resident macrophages could contribute to age-related changes, such as impaired tissue repair.

The liver comprises the largest population of resident macrophages in the body. These Kupffer cells (KCs) sit in the liver fenestrated sinusoidal endothelial cells, where they are in close contact with bloodstream circulation. In the aged rodent liver, alteration of the microvasculature^17^ activation of stromal cells and decline in hepatocyte function have been observed, along with general inflammation. Moreover, a higher prevalence of inflammation related chronic liver diseases (cirrhosis, non-alcoholic steatohepatitis) has been observed in aged individuals^18^. Recently, the Covid-19 pandemic, which disproportionately affect elderlies and people with chronic metabolic pathologies, has highlighted the need to disentangle the interplay between inflammation, ageing and tissue repair. Since KCs are established key players of the inflammation and tissue repair processes, a better understanding of their maintenance and functions in physiological ageing will thus prove critical for the treatment of age-related pathologies.

Because embryo-derived macrophages are usually thought to be replaced by HSC-derived cells that can adopt a similar phenotype, in particular after acute macrophage depletion ^11,12,19^, we expected to observe an increased contribution of HSC-derived cells to tissue resident macrophages with age without changes in their overall tissue density. Here we show that resident macrophage density decreases 2-fold during ageing in the liver and other solid organs, and this resident macrophage loss is associated with liver tissue dysfunction. The net contribution of HSC-derived cells to resident macrophage populations is constant over time and no compensatory recruitment of monocyte-derived cells was observed. This loss was specific to KCs, when compared to other innate immune cells present in the liver and was not due to proliferative exhaustion. Furthermore, sustained inflammation is sufficient to drive resident macrophage loss while impaired inflammation sensing rescues the age-associated phenotype. Our study demonstrates that increased niche availability, defined here as available space due to a decrease in macrophage density in combination with available CSF1 growth factor presentation by Hepatic Stellate cells, is not sufficient to elicit the recruitment and local differentiation of HSC-derived cells. These data support a model where inflammation triggers the net loss of resident macrophages by modifying the homeostatic proliferation/death balance, thereby leading to a cumulative loss of resident macrophages over time without compensation from HSC-derived cells. Moreover, preventing liver resident macrophage sparseness improved age-associated tissue dysfunction. Thus, further understanding the factors regulating resident macrophage homeostasis and the signals mediating cell recruitment in the context of ageing will lead to new strategies to improve aged tissue homeostasis and repair.

## Results

### Kupffer cell numbers are reduced with Age

We scored phenotypic resident macrophages in the liver, named Kupffer cells (KCs), defined here by the expression of the cell-surface markers F4/80, CD11b and Tim4 (<u>Figure S1A</u>). Four age groups were studied: Young mice (2-4 months-old, mO) were compared with three groups of aged mice (16-18mO, 20-25mO and 29-30mO). Interestingly, phenotypic resident macrophage density started to decrease at 16-18 months, with an average 2-fold-reduction at 20-25 months (Figure 1A). These results were confirmed by counting F4/80+ macrophages *in situ* in liver sections (Figure S1B). The reduction in density of macrophages with age was specific to resident macrophages, as the density of other liver innate immune populations, such as neutrophils and MHC-II^+^ CD11b^high^ F4/80^-/low^ cells, were increased at 20-25 months and 29-30mO, respectively (Figure 1B and <u>Figure S1C</u>). Given the increased proportion of neutrophils among circulating cells in peripheral blood at 20-25 months of age (Figure 1C), the increase in liver neutrophil density could be the reflection of increased circulating cells and represent an increase of neutrophils within the liver sinusoids. To better assess neutrophil location (circulation *vs* intra-tissue), we performed an intravenous injection of anti-CD45 antibody. We evaluated the proportion of liver monocytes and neutrophils that were labelled by this antibody and normalised them to the labelling observed on their circulating counterparts. We found that in young and aged animals, Ly-6C^hi^ monocytes were equally labelled in the blood and in the liver. In contrast, neutrophil labelling was decreased with age in the liver when compared to circulating cells. This suggests a transmigration of neutrophils from the liver circulation to the liver parenchyma (<u>Figure S1D</u>). Histological examination confirmed the increase in tissue MPO^+^ neutrophils (<u>Figure S1E</u>) and demonstrated that the proportion of neutrophils that were intra-tissular (not located within CD31^+^ blood sinusoid vessels) was increased in aged livers (<u>Figure S1F</u>). Collectively, these results suggest that the density of neutrophils within the liver could be a used as read-out of the low-grade sustained inflammation that is postulated to occur in aged tissues.

**Figure 1:**
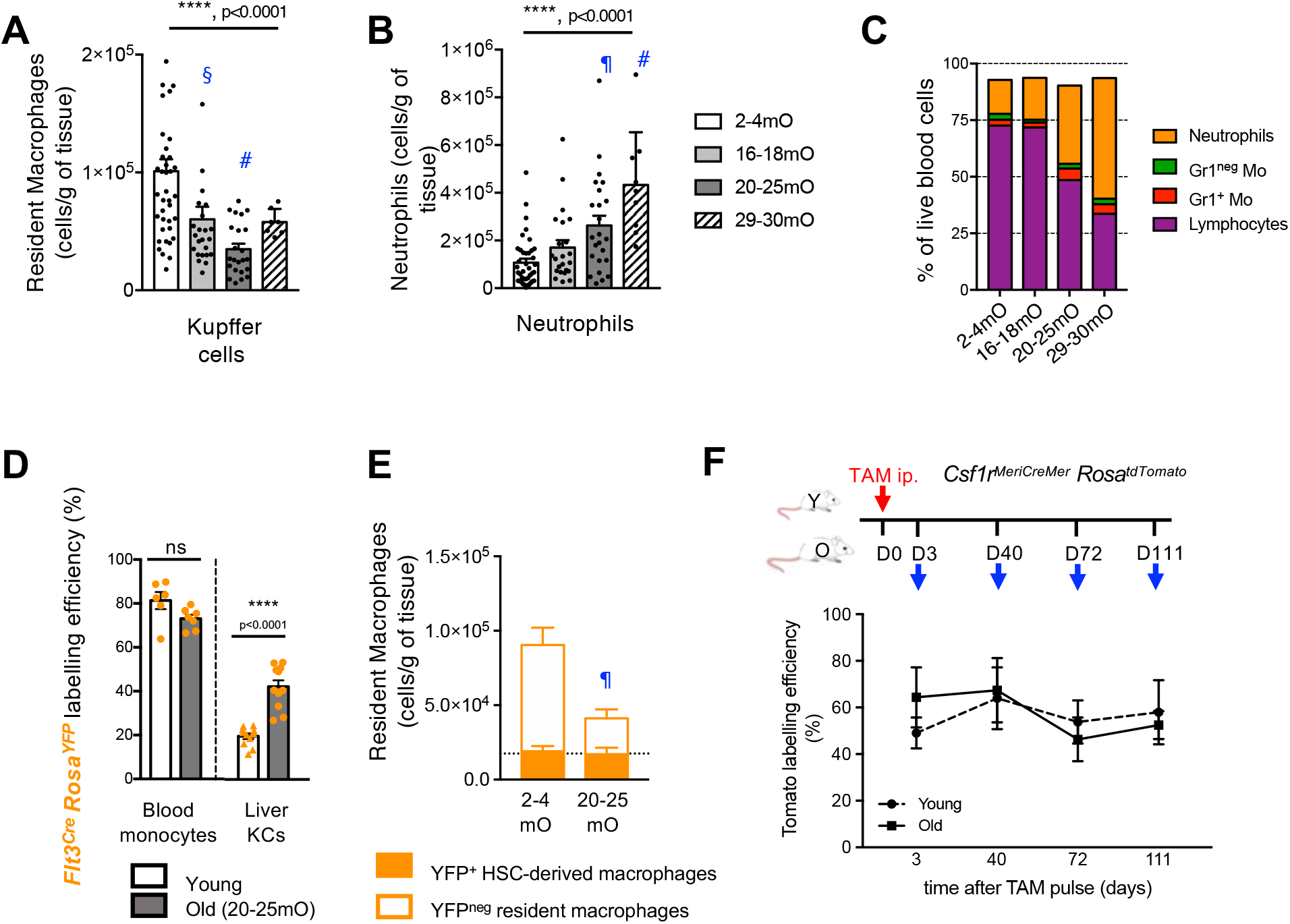
Loss of Kupffer cells with age is not compensated by HSC-derived cells. **A**, Number of liver resident macrophages (Kupffer cells) per gram (g) of tissue and **B**, Number of neutrophils per g of liver from Young 2-4 month-old (mO) (n=40), and Old 16-18mO (n=23), 20-25mO (n=24) and 29-30mO (n=9) animals. Mean ± s.e.m; one-way ANOVA (Kruskal Wallis) and post-test Dunn multiple comparison test against Young group: #, p<0.0001; ¶, p<0.001; §, p<0.01. **C**, Frequency of neutrophils, Gr1^-^ and Gr1^+^ monocytes (Mo) and lymphocytes in blood from 2-4 mO (n=36), 16-18mO (n=17), 20-25mO (n=29) and 29-30mO (n=12) animals. **D**, YFP labelling efficiency in blood monocytes and liver resident macrophages (Kupffer cells, KC) in Young (2-4 mO, n=10) and Old (20-25 mO, n=8) *Flt3*^*Cre*^ *Rosa*^*YFP*^ mice. Mean ± s.e.m; Mann-Whitney test. **E**, Number of (*Flt3*^*Cre*^)-YFP^+^ and (*Flt3*^*Cre*^)-YFP^neg^ resident macrophages per g of liver. Dashed line represents the mean density of (*Flt3*^*Cre*^)-YFP^+^ resident macrophages, mean ± s.e.m; Young (2-4mO, n=10) and Old (20-25mO, n=8). Mean ± s.e.m; Dunn multiple comparison test between Young and Old (*Flt3*^*Cre*^)-YFP^neg^ resident macrophages: ‡, p<0.05. **F**, Pulse labelling of *Csf1r*-expressing cells in Young (2-4 mO) and Old (24-31 mO) *Csf1r*^*MeriCreMer*^ *Rosa*^*tdTomato*^ mice by a single injection of tamoxifen (TAM) and analysis 3 days (D3), 40 days (D40), 72 days (D72) and 111 days (D111) after. Tomato labelling efficiency in liver KCs, Mean ± s.e.m; two-way ANOVA analysis; n=3 per age group and per timepoint.

### Kupffer cells are not replaced by HSC-derived cells during Ageing

Recent literature suggested that the contribution of HSC-derived cells would increase over time, and the reduction in the total number of macrophages does not preclude this possibility. We thus sought to investigate the contribution of HSC-derived cells to phenotypic resident macrophages during ageing by using the *Flt3*^*Cre*^ *Rosa*^*YFP*^ fate mapping model^20,21^. HSC-derived cells are efficiently labelled in *Flt3*^*Cre*^ *Rosa*^*YFP*^ mice, with 80% of circulating monocytes been YFP^+^. The proportion of HSC-derived (*Flt3*^*Cre*^)-YFP^+^ cells among resident macrophages increased with time in the liver (Figure 1D). However, because the population size changed over time as demonstrated in Figure 1A, we examined the density of (*Flt3*^*Cre*^)-YFP^+^ and -YFP^neg^ resident macrophages. While there was no change in HSC-derived (*Flt3*^*Cre*^)-YFP^+^ cells, we observed a specific loss of (*Flt3*^*Cre*^)-YFP^neg^ cells (2-fold reduction at 20-25mO when compared to Young) (Figure 1E). Thus, the observed reduction of resident macrophage density was specific to HSC-independent macrophages and was not compensated by increased recruitment and/or differentiation of HSC-derived cells. This result was confirmed using a complementary *Csf1r*^*MeriCreMer*^ *Rosa*^*tdTomato*^ approach, where macrophages expressing the inducible Cre recombinase (MeriCreMer) under the control of the macrophage growth factor receptor (Colony stimulating factor receptor 1, *Csf1r*) promoter were pulse-labelled in Young and Old mice. Pulse-labelling of macrophages in Young and Old *Csf1r*^*MeriCreMer*^ *Rosa*^*tdTomato*^ mice led to efficient recombination (evidenced by a Tomato labelling efficiency of 60%) in liver resident macrophages three days after a single dose of tamoxifen (Figure 1F). Contribution of (non-labelled) HSC-derived cells to the resident macrophage population would lead to a decline in Tomato labelling over time. There was no significant difference in macrophage Tomato labelling between day 3 and day 111 post tamoxifen injection between Old and Young, suggesting that macrophage self-maintenance or half-life in aged livers was not affected when compared to young livers (Figure 1F). Collectively, this indicates that there was no switch in the ontogeny of liver resident macrophages with ageing. In contrast to listeria infection, whole body-irradiation or DT-mediated ablation of KCs ^11,12,19,22^, HSC-derived cells were either not recruited or failed to differentiate into tissue macrophages when liver macrophage density was reduced with age.

### Kupffer cells display minor changes with Age

We next investigated whether liver resident macrophages (KCs) were functionally affected with ageing. A functional feature of liver KCs is their ability to phagocyte large particles directly from the bloodstream, due to their localization within the liver sinusoids. KC phagocytosis of 2µm PE-latex beads was unaffected in Old animals when compared to Young controls (Figure 2A). However, we observed morphological changes in old liver resident macrophages, with Old KCs larger in size and they had increased granularity (<u>Figure S2A and B</u>). Old KCs also had increased cell-surface expression of the macrophage marker F4/80 and the myeloid integrin CD11b, as measured by the median fluorescent intensity. To take into account the increased cell size, F4/80 and CD11b expression were normalized to cell size (FSC-A). While normalized F4/80 expression was not changed between Young and Old, the increase in CD11b expression could not be explained alone by the increased cell size (<u>Figure S2C</u>). An exploration of the Tabula Muris Senis Consortium RNA sequencing repository ^23^ showed an increase in the number of reads in KCs, a feature traditionally associated with increased cell size. This increase in reads and expressed genes with age was specific to KCs, as hepatocytes were unaffected (Figure S2D).

**Figure 2:**
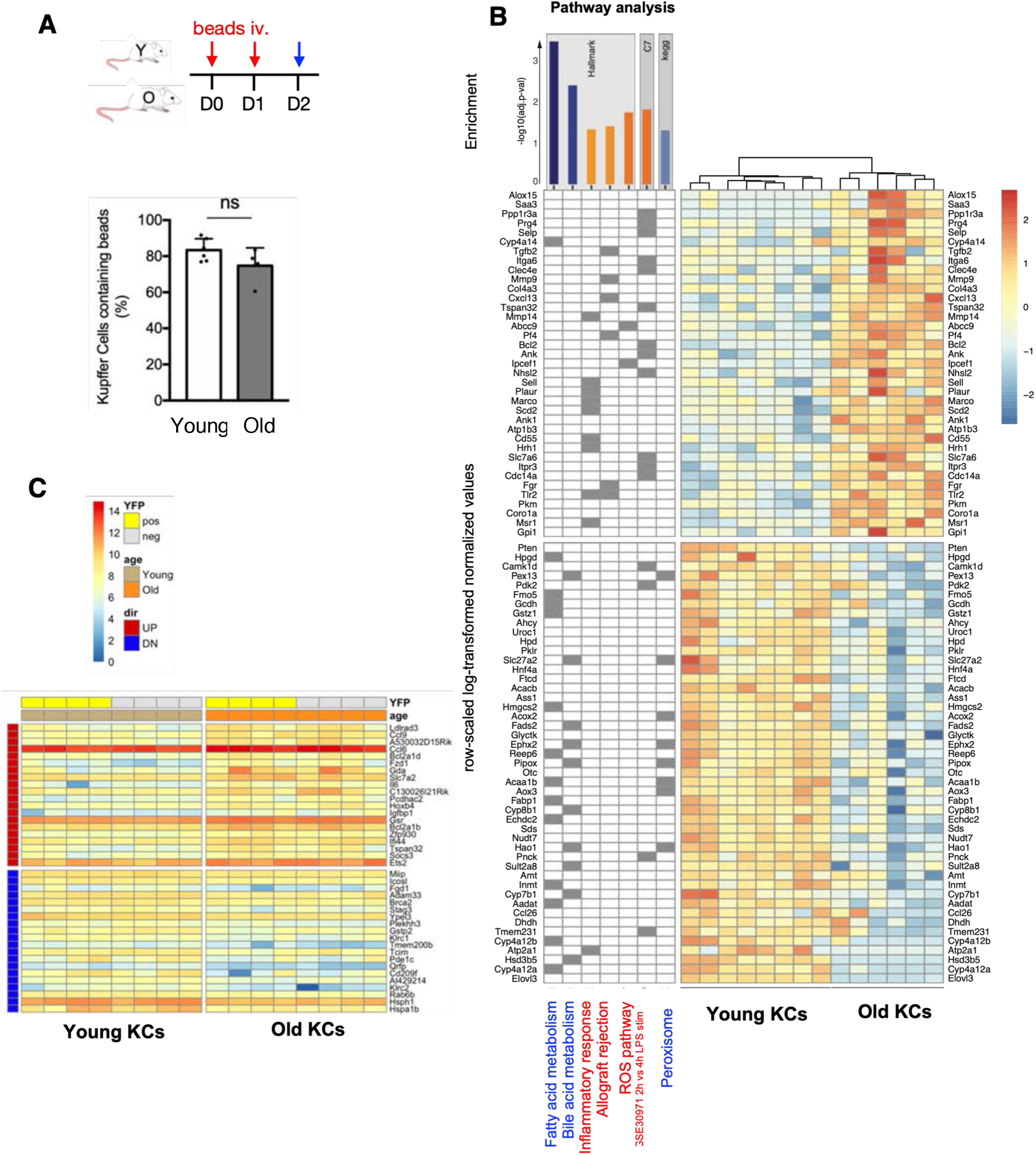
Kupffer cells show few intrinsic changes with age. **A**, Analysis of in vivo phagocytosis of 2µm beads injected intravenously (iv., red arrows). Frequency of Kupffer cells containing beads 24h after the last injection from Young (3 mO; n=6) and Old (19 mO; n=4) animals. **B**. RNA-sequencing analysis of Young (4mO) and Old (22-23mO) liver KCs. Right panel : Heat map of the up- or down-regulated genes between old and young KCs (log-transformed normalized values scaled by rows). Dendrogram shows the hierarchical clustering using the Euclidean distance between samples. Left panel : binary matrix showing the genes affiliation to the gene collections found enriched either in the young KCs (lower panel) or in the old KCs (upper panel). Top panel : Barplot displaying the adjusted p-value (-log10-transformed) of the gene set enrichment analysis (GSEA). Blue, and orange bars for gene collections upregulated in young or old KCs, respectively. **C**, RNA sequencing analysis of sorted Young and Old KCs from *Flt3*^*Cre*^ *Rosa*^*YFP*^ mice, sorted based on YFP expression: YFP^+^ (yellow) and YFP^neg^ (grey) cells. Heat map shows up- (red) and down- (blue) regulated genes between young and old KCs.

Whole transcriptomic analysis of Old (22-23 mO) versus Young (2-4 mO) KCs showed only 37 up-regulated genes and 47 down-regulated genes (Figure 2B). Gene set enrichment analysis revealed an up regulation for pathways related to inflammatory response, leukocyte migration and LPS-stimulated macrophages in old KCs, while pathways for peroxisome, fatty acid and bile acid metabolism were significantly up regulated in young KCs (Figure 2B). We further compared KCs in young and old *Flt3*^*Cre*^ *Rosa*^*YFP*^ mice; and focused on (*Flt3*^*Cre*^)-YFP^neg^ and (*Flt3*^*Cre*^)-YFP^+^ Kupffer Cells. Here too, we found only minimal differences (Figure 2C), in line with the importance of tissue environment in instructing resident macrophage phenotype and transcriptome ^24^. Thus, age minimally changed the liver resident macrophage transcriptome (as recently show for aged neural stem cells^25^) and old KCs appeared to retain their self-maintenance and phagocytic abilities.

### Kupffer Cell proliferation is not altered with Age

Resident macrophages are maintained in adult tissues by local proliferation^26–31^ and we could not detect any changes in the proliferation of liver macrophages between Young and Old animals (Figure 3A). When distinguishing in *Flt3*^*Cre*^ *Rosa*^*YFP*^ KCs between different ontogenies, proliferation of HSC-independent (*Flt3*^*Cre*^)*-*YFP^neg^ KCs was unaffected in Old, as measured by Ki67 immunostaining *in situ* (Figure 3A) or by flow cytometry after EdU incorporation (three consecutive injections) (Figure 3B). Further, EdU incorporation in HSC-derived (*Flt3*^*Cre*^)*-*YFP^+^ macrophages was also comparable between young and Old. This suggested that proliferative exhaustion was not responsible for the reduction in resident macrophage density. Although basal levels of proliferation were unaffected, old KCs may fail to mount a proliferative burst in response to injury or challenge. A classical model to induce a proliferative burst in macrophages is withdrawal of CSF1R inhibition. CSF1 is the main growth factor supporting the maintenance and proliferation of most adult macrophages, including Kupffer Cells. We targeted the CSF1 receptor (CSF1R) with the blocking compound Plexikkon 5622 (PLX)^32^ and inhibited CSF1R signalling for three days, which led to KC depletion in both young and aged livers (Figure 3C). After CSF1R inhibition withdrawal, mice were allowed to recover for 9 days and KC proliferation was measured by a single EdU pulse two hours before analysis. Young and old KCs had similar proliferation levels at all timepoints (Figure 3C). Importantly, Old KCs displayed a two-fold increase in their proliferation rate in the recovery phase when compared to control, demonstrating that age did not affect KC proliferative potential nor the ability of KC to respond to CSF1 stimulation (Figure 3C). Further, KC proliferative potential was highlighted by the replenishment of the KC compartment after depletion at a similar rate between young and Old (Figure 3B), suggesting similar proliferative potential. Finally, analysis of YFP labelling efficiency in *Flt3*^*Cre*^ *Rosa*^*YFP*^ after PLX recovery demonstrated no significant increase in the input from bone marrow progenitors or circulating cells to the repopulated KC pool (Figure 3D).

**Figure 3:**
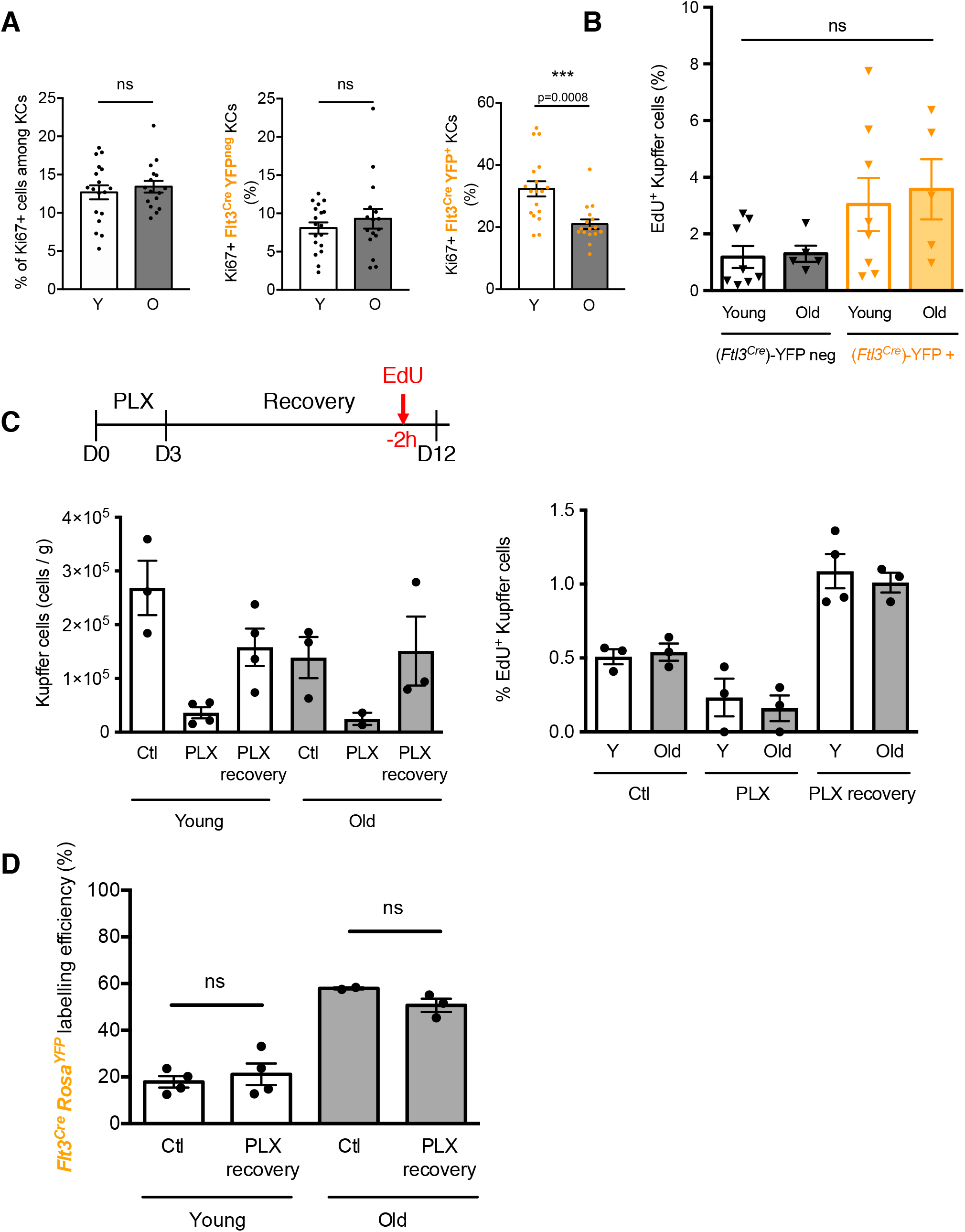
Kupffer cell proliferation is not altered with age. **A**, Frequencies of proliferating Ki67^+^ F4/80^+^ Kupffer cells (KCs), Ki67^+^ F4/80^+^ *Flt3*^*Cre*^ YFP^neg^ (HSC-independent), and Ki67^+^ F4/80^+^ *Flt3*^*Cre*^ YFP^pos^ (HSC-derived) KCs in liver cryo-sections from Young (2-4 mO) and Old (20-25 mO) *Flt3*^*Cre*^ *Rosa*^*YFP*^ mice. Mean ± s.e.m; Mann-Whitney test; n=3 per age group. **B**, Frequencies of proliferating EdU+ KCs in Young (2-4 mO) and Old (20-25 mO) *Flt3*^*Cre*^ *Rosa*^*YFP*^ mice. Mean ± s.e.m; Mann-Whitney test; n= 8 and 5 for young and old, respectively. **C**, Number of liver resident macrophages (Kupffer cells) per gram of liver (Left) and Frequencies of proliferating EdU+ KCs (right) in Young and Old mice in control (CTL), after PLX treatment and after PLX treatment with 9-day recovery. Left: Two-Way ANOVA: significant effect of the treatment factor (p= 0.0037); no significant effect of the age factor (p=0.1633). Right: Two-Way ANOVA: significant effect of PLX treatment (p<0.0001); no significant effect of age (p=0.7360). **D**, YFP labelling efficiency in Young (2-4 mO) and Old (20-25 mO) *Flt3*^*Cre*^ *Rosa*^*YFP*^ mice, in CTL or after PLX treatment and recovery (PLX recovery).

### Most tissue resident macrophages are not replaced by HSC-derived cells

The age-associated reduction of resident macrophage numbers was observed in other major solid organs, with the exception of the kidney (<u>Figure S3A</u>). This reduction was also specific to resident macrophages, as other resident immune populations (dendritic epidermal T cells, DETCs) and other myeloid populations were stable until at least 20-25mO (e.g. F4/80^bright^ CD64^+^ CD11b^+^ interstitial lung macrophages, Siglec-F^+^ F4/80^low^ CD11b^+^ lung eosinophils and spleen Ly-6C^+^ CD11b^high^ F4/80^-/low^ cells) (not shown). There was also no age-associated switch in the ontogeny of resident macrophages in the epidermis (Langerhans cells), lung (alveolar macrophages), spleen (red pulp macrophages), and brain (microglia) with ageing that would allow the compensation of resident macrophage loss (<u>Figure S3B</u>). In contrast, the number of (*Flt3*^*Cre*^)-YFP^+^ macrophages seemed increased at 20-25mO in the kidney (<u>Figure S3B</u>). It thus appeared that kidney resident macrophage density was in part maintained during ageing by an increased recruitment and differentiation of HSC-derived cells. Altogether, the density of most resident macrophages, with the exception of kidney macrophages, declined with age without compensation from HSC-derived cells.

### Poly I:C-induced repeated inflammation induces KC cell death, mimicking the ageing phenotype

Since proliferative capacity was not impaired in aged KC, we next investigated whether increased cell death could be responsible for the observed KC loss in ageing. In aged livers, we observed a 5-fold increase in the proportion of Tunel positive Kupffer Cells compared to young, suggesting that gradual and cumulative KC cell death could be driving the aged phenotype (Figure 4A). Age is associated with low-grade sustained inflammation (reviewed in ^34^) and we found an up-regulation of pro-inflammatory pathways in Old KCs (Figure 2B), as recently reported also in aged microglia cells^25^. Since inflammation has been shown to drive macrophage cell death^33^, we thus investigated whether sustained inflammation with a viral mimic, Poly(I:C), could drive resident macrophage loss. First, acute Poly(I:C) challenge did not affect resident macrophage density in the liver, nor any other organ studied (<u>Figure S4</u>). Sustained stimulation was performed by injecting every other day Poly(I:C) (5mg/kg) or saline for a month (Figure 4B). Two weeks after the end of the Poly(I:C) treatment, resident macrophage density was decreased 2-fold in the liver (Figure 4C). This loss of KC was not transient, as it was still observed one month later (6 weeks post last injection, pli). In order to investigate whether sustained inflammation was driving the specific loss of HSC-independent resident macrophages as shown during ageing, we next examined *Flt3*^*Cre*^ *Rosa*^*YFP*^ mice subjected to sustained Poly(I:C) and analysed 2 weeks and 6 weeks pli. As observed during ageing, there was a specific loss of HSC-independent (*Flt3*^*Cre*^)-YFP^neg^ cells in liver while HSC-derived (*Flt3*^*Cre*^)-YFP^+^ cells remained at comparable levels (Figure 4D). Sustained Poly(I:C) only elicited a transient neutrophil density increase 48h pli in the liver that resolved by 2 weeks pli (Figure 4C). Thus, sustained inflammation with Poly(I:C) did not fully recapitulate the ageing phenotype in regard to neutrophils. Further, these results evidenced that inflammation stimuli was not sufficient to induce monocyte recruitment and differentiation into macrophages, in contrast to what is observed after ‘total’ depletion of KC ^11,12^. All tissues examined showed a similar decrease of resident macrophage density after sustained Poly (I:C) (1.2-fold decrease in epidermis, 1.7-fold in spleen, 2.3-fold in lung and 2.6-fold in kidney), except in the brain (<u>Figure S5A</u>). Further, Tunel staining on liver sections showed an increase in the proportion of Tunel+ positive Kupffer cells two weeks post Poly (I:C) treatment, a phenotype that was maintained 6 weeks post treatment (Figure 4E). As in aged mice, sustained inflammation induced KC cell death that led to a decreased KC density, without compensation (by either monocyte recruitment or local proliferation).

**Figure 4:**
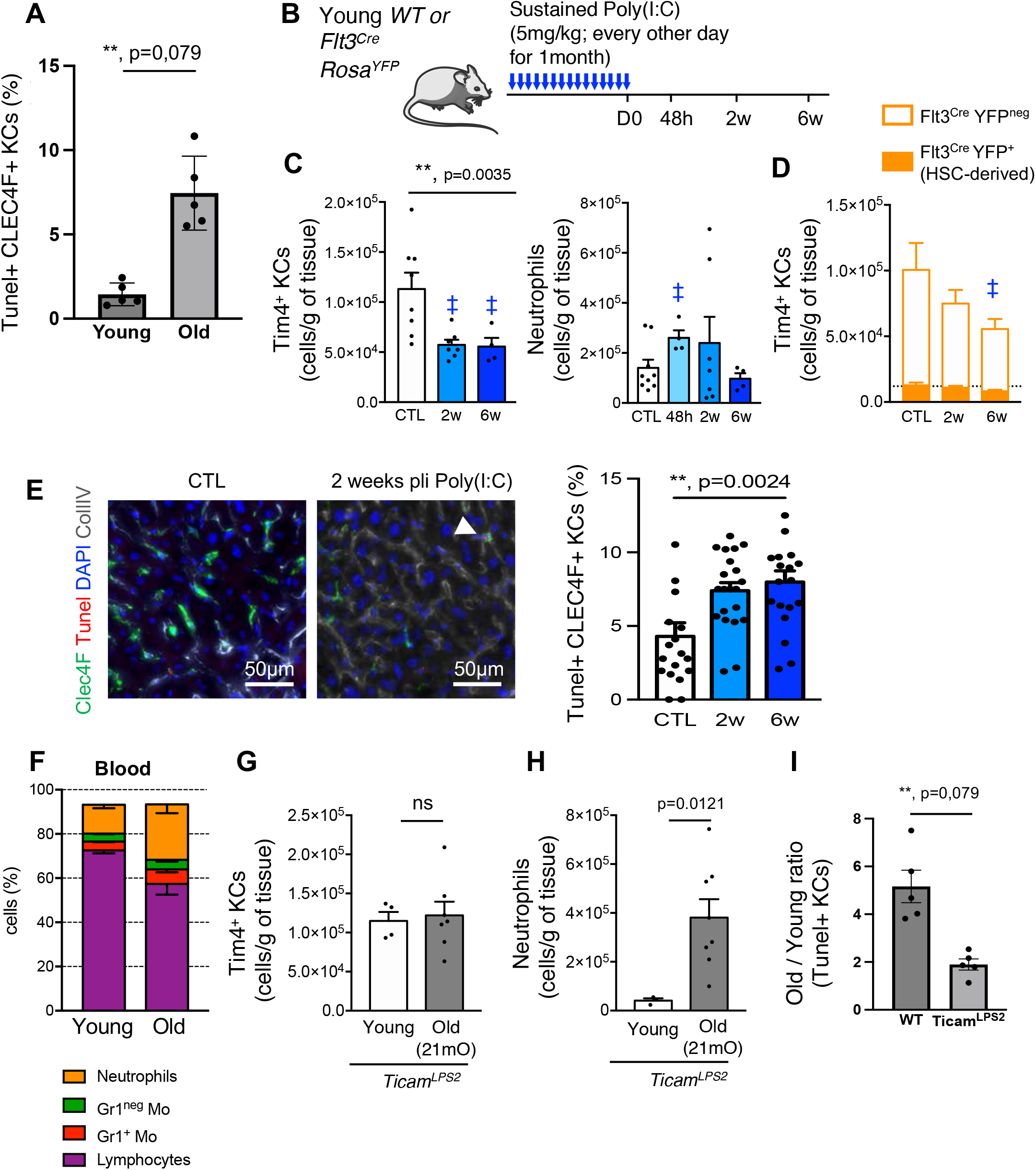
Sustained inflammation mimics the age-related KC phenotype while KC cell death and loss is attenuated in inflammation sensing deficient *Ticam*^*LPS2*^ mice. **A**, Frequencies of apoptotic Tunel+ F4/80^+^ CLEC4F+ Kupffer cells (KCs), in liver cryo-sections from Young (2-4 mO) and Old (20-25 mO) *Flt3*^*Cre*^ *Rosa*^*YFP*^ mice. Mean ± s.e.m; Mann-Whitney test; n=5 per age group. **B**, Experimental protocol for sustained poly I:C injections and **C**, Number of liver resident macrophages (Kupffer cells) per gram of liver from saline (CTL), 2w and 6w after sustained Poly(I:C) injections in Young adult mice. Mean ± s.e.m; Dunn multiple comparison test against saline control group: ‡, p<0.05; saline controls, n=9-11; Poly(I:C) 2w, n=8 and Poly(I:C) 6w, n=4. **D**, Number of Number of liver resident macrophages (Kupffer cells) per gram of liver from saline (CTL), 2w and 6w after sustained Poly(I:C) injections in Young adult *Flt3*^*Cre*^ *Rosa*^*YFP*^ mice. Mean ± s.e.m; Dunn multiple comparison test against saline control group. **E**, Representative liver cryo-sections of saline (CTL) or 2w after sustained Poly (I:C) injection in young adult mice, stained with CLEC4F (green), Coll IV (grey), DAPI (blue) and Tunel (red) and quantification of the frequencies of apoptotic Tunel+ F4/80+ CLEC4F+ Kupffer cells (arrowhead). Scale bar represents 50µm. **F**, Frequency of neutrophils (orange), Gr1^-^ and Gr1^+^ monocytes (green and red, respectively) and lymphocytes (purple) in blood from Young (5 mO, n=5) and Old (21mO, n=8) *Ticam*^*Lps2*^ mutant animals. **G**, Number of neutrophils and **H**, Number of KCs per g of liver from Young (5 mO, n=5) and Old (21mO, n=8) *Ticam*^*Lps2*^ mutant animals. Mean ± s.e.m; Mann-Whitney test. **I**, Ratio of the proportion of Old Tunel+ KCs compared to the proportion of Young Tunel+ KCs in WT and *Ticam*^*LPS2*^ mice. Mann Whitney test; ****: p<0,0001.

### KC age-related phenotype is attenuated in immune sensing *Ticam* ^*LPS2*^ deficient mice

To investigate whether systemic gradual, cumulative inflammation could be responsible for the resident macrophage loss observed during ageing, we next analysed mutant mice where inflammation sensing is defective, but not abolished. We took advantage of *Ticam*^*Lps2*^ mutant mice that have a point mutation in the signalling protein TICAM/TRIF that is downstream of TLR signalling^35^. *Ticam*^*Lps2*^ mutant mice lack cytokine responses to Poly(I:C) and have severely impaired responses to LPS. 21-month-old *Ticam*^*Lps2*^ mutant mice had increased blood and liver neutrophils when compared to young mutant mice (Figure 4 F and H), as observed in *wild-type* (WT) ageing. However, *Ticam*^*Lps2*^ mutation rescued the ageing phenotype observed in KCs (Figure 4G). The same was observed in all tissues, as there were no differences in resident macrophage density between Young and Old *Ticam*^*Lps2*^ mutant mice (<u>Figure S6</u>). Moreover, the 5-fold increase in dying (Tunel+) KC in old WT animals was attenuated in old *Ticam*^*Lps2*^ KCs (Figure 4I).

Therefore, impaired inflammation signalling specifically rescued the age-associated resident macrophage loss in all tissues examined. In combination with the sustained inflammation challenge, our data shows that cumulative inflammation drives resident macrophage cell death, which in turn leads to the age-associated reduction in resident macrophage density. Further, our results indicate that the decrease in macrophage density is not required to induce the accumulation of tissue neutrophils and conversely, that the neutrophil infiltration is not sufficient to induce macrophage loss.

### Age-induced KC loss is not associated with changes in trophic factors availability

The loss of resident macrophages, combined with the inability of HSC-derived cells to infiltrate the liver tissue and replace them, may reflect a change in the liver environment that renders it less supportive of macrophage survival, proliferation and/or differentiation. Stellate cells, hepatocytes and endothelial cells have been proposed as the key actors responsible for Kupffer cell identity and maintenance^12^. In particular, Hepatic Stellate Cells have been shown to support survival and proliferation of KCs by producing Interleukin 34 (IL34) and Colony Stimulating Factor 1 (CSF-1), the two ligands of CSF1R. We thus investigated whether changes in hepatic stellate cells with age could contribute to the KC phenotype. First, we quantified the number of Desmin+ hepatic stellate cells and found no significant change that could correlate with the loss of KCs with age (Figure 5A and B). We then investigated whether aged stellate cells had impaired CSF1 and/or IL34 presentation at their cell surface (Figure 5A). In WT mice, the proportion of CSF1-expressing stellate cells was increased (Figure 5B), which could reflect an increased production of CSF1 to help sustain a dwindling pool of macrophages. Ageing did not differentially affect *Ticam*^*Lps2*^ stellate cells compared to WT, suggesting that this increase in CSF1 production may not be solely due to KC decreased density (Figure 5B). In young mice that received repeated injections of Poly(I:C), the number of Desmin+ cells remained stable up to 6 weeks post last injection, but the proportion of CSF1 and IL34 producing cells was reduced at 2weeks (Figure 5C) before returning to steady-state levels at 6 weeks post last injection. This suggests a specific effect of Poly (I:C) on stellate cells. Altogether, diminished availability of CSF1 and of hepatic stellate cells is not responsible for the lack of monocyte-derived macrophage engraftment in old livers.

**Figure 5:**
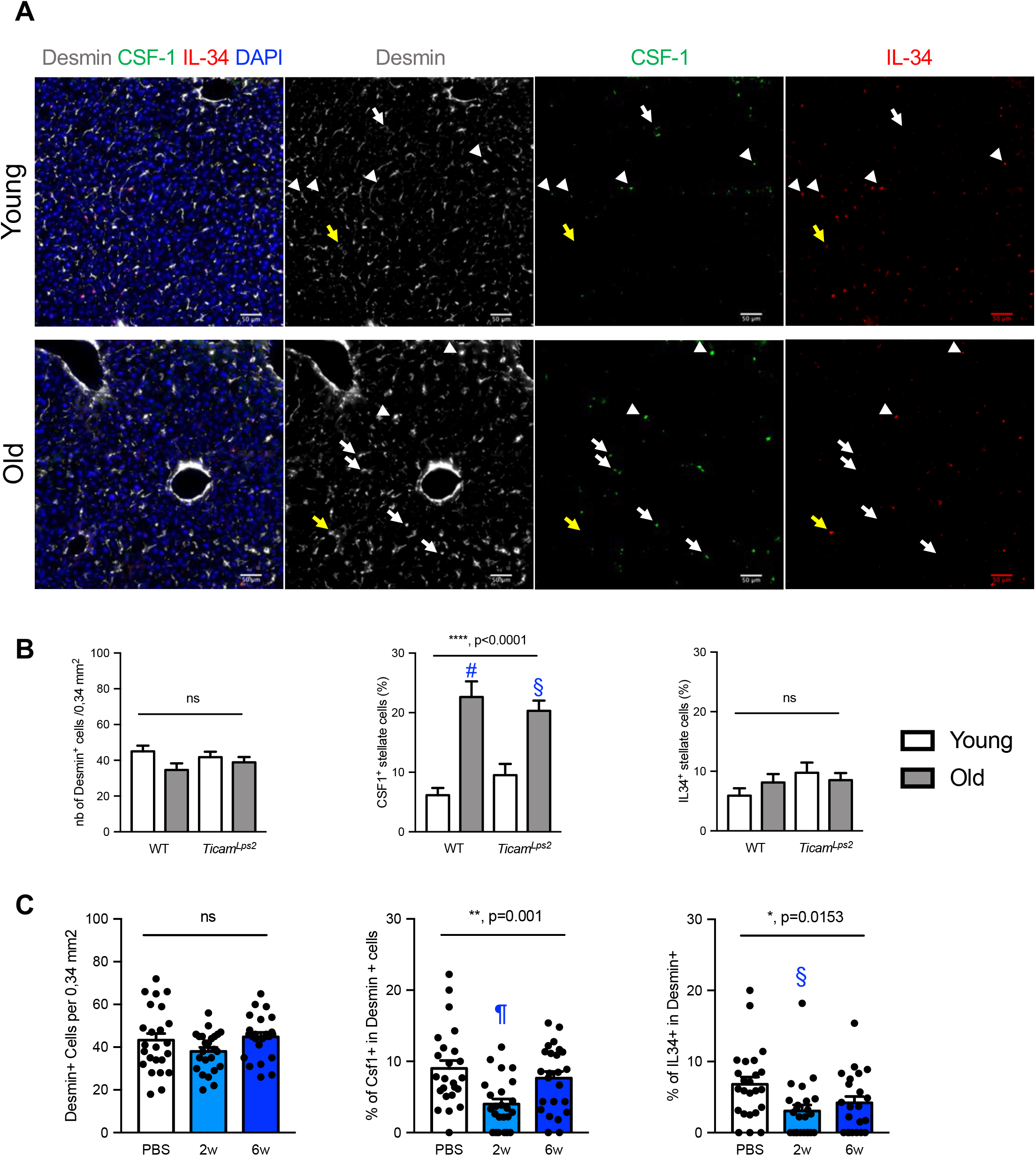
Changes in trophic factors availability do not explain KC loss with age. **A**, Representative liver cryo-sections from Young (2-4 Mo) and Old (21-24 Mo) mice stained with Desmin (grey), CSF1 (green), IL34 (red) and Dapi (blue). Scale bar represents 50µm; white and yellow arrows point to CSF1+ Desmin+ and IL34+ Desmin+ hepatic stellate cells, respectively. **B**, Proportion of Desmin positive (left panel), CSF1+ Desmin+ (middle panel) and IL34+ Desmin+ (right panel) in liver sections from Young and Old WT and *Ticam*^*Lps2*^ mice. n=5 per age. **C**, Proportion of Desmin positive (left panel), CSF1+ Desmin+ (middle panel) and IL34+ Desmin+ (right panel) in liver sections from saline (PBS), 2w and 6w after sustained Poly(I:C) injections in Young adult mice. Mean ± sem. One-way ANOVA (Kruskal Walis): #, p<0.0001; ¶, p<0.001; §, p<0.01.

### Hallmarks of liver aging are attenuated when inflammation sensing is impaired

Finally, to characterize the functional relevance of resident macrophage density during ageing, we investigated key features of liver age-associated dysfunction: increased lipid content and senescent cell accumulation, in both WT and *Ticam*^*Lps2*^ mutants. As reported previously^36^, we observed a massive increase in senescent cell and lipid contents accumulation in the aged liver (Figure 6). Importantly, when macrophage density is maintained in old *Ticam*^*Lps2*^ mice, senescent cell (Figure 6A-B) and lipid droplet accumulation with age was attenuated (Figure 6C). Thus, maintaining a youthful resident macrophage density seems to be critical to prevent key features of liver dysfunction with age.

**Figure 6:**
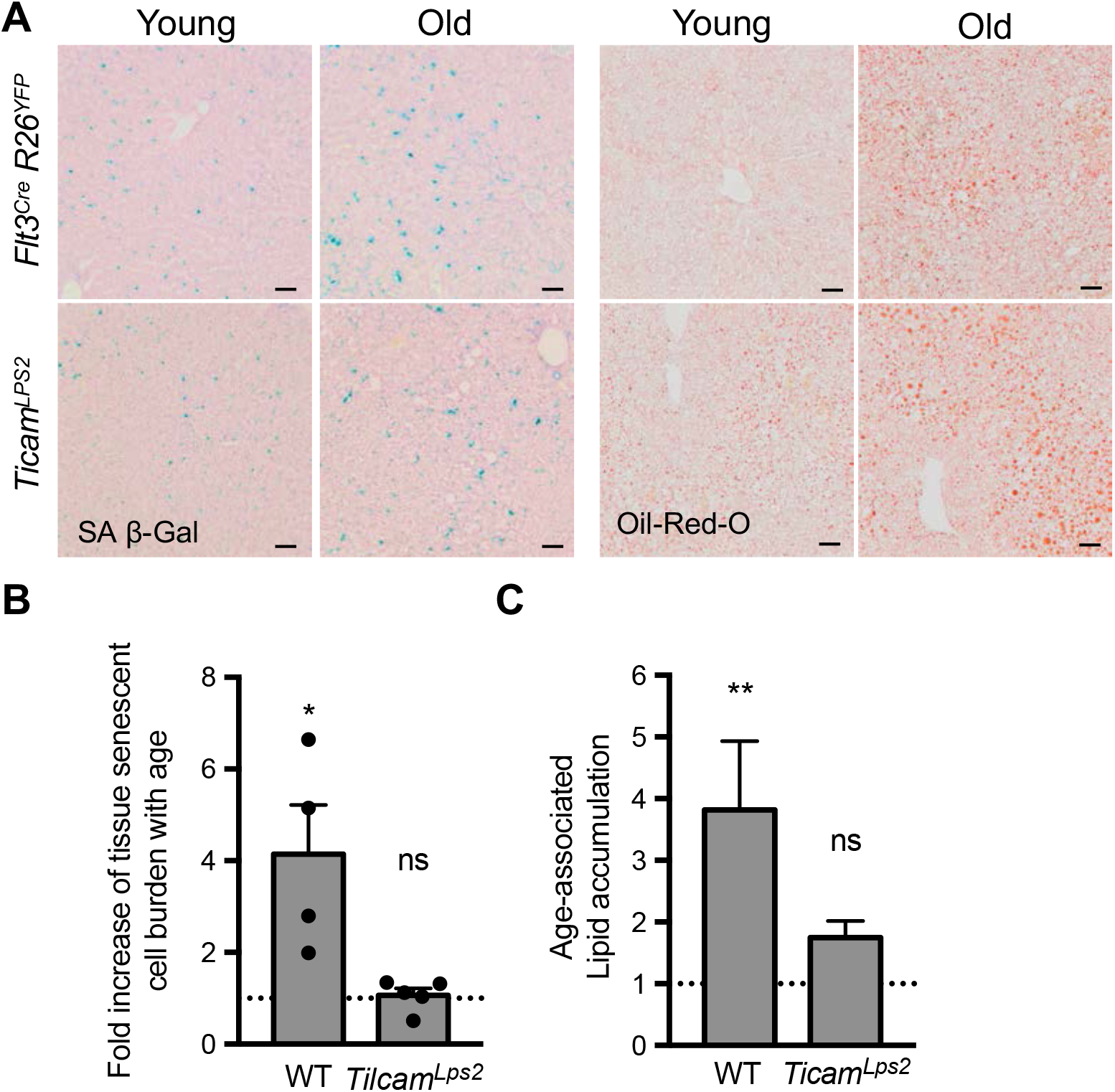
Old KC density correlates with hallmarks of liver aging: increased lipid and senescent cell accumulation. **A**, Representative images of SA β-Gal and Oil-Red-O stained liver sections from Young and Old *Flt3*^*Cre*^ *Rosa*^*YFP*^ mice and *Ticam*^*Lps2*^ mice. Bar: 50 µm **B**, Fold change (old /young) in tissue senescent cell burden as measured by the SA β-Gal positive area in liver sections from Young and Old *Flt3*^*Cre*^ *Rosa*^*YFP*^ mice and *Ticam*^*Lps2*^ mice. Mean ± s.e.m. **C**, Fold change (old/young) or lipid accumulation measured by the Oil-Red-O positive area in liver sections from Young and Old *Flt3*^*Cre*^ *Rosa*^*YFP*^ mice and *Ticam*^*Lps2*^ mice. Mean ± s.e.m *: t-test, *:p<0,05; **:p<0,01 compared to young of the same genetic background.

## Discussion

Collectively, our findings demonstrate that there is a specific loss of HSC-independent tissue resident macrophages during ageing in the liver and most other tissues examined. Loss of HSC-independent resident macrophages was not explained by proliferative exhaustion and was not compensated by increased local recruitment and/or differentiation of HSC-derived cells. Instead, we found that sustained inflammation is sufficient to induce the age-associate macrophage phenotype in the liver, while reduced inflammation sensing during ageing abolished specifically the resident macrophage loss.

These data support a model where inflammation specifically impacts HSC-independent resident macrophages by triggering an imbalance in their proliferation/death homeostasis, thereby leading to their cumulative loss over time without compensation from HSC-derived cells. While HSC-derived cells can compensate for the loss of HSC-independent resident macrophages in very specific experimental paradigms, as recently illustrated for the liver^19,22^, we did not observe this phenomenon during physiological ageing or sustained inflammation in the liver, which raises the question of why HSC-derived cells fail to infiltrate and differentiate into tissue macrophages in aged tissues. Among the different liver cells that provide supportive signals to KCs, liver sinusoid cells undergo profound changes and remodelling during ageing, in a process called pseudo-capillarisation, and could play a significant but yet under-recognized role in this process. Thus, further work is needed to identify the factors regulating resident macrophage homeostasis and the signals mediating cell recruitment and survival in different physio-pathological conditions.

It has been theorized that accumulation of gradual sub-clinical tissue damage (in response to sterile inflammation or infection) increases the burden of tissue repair with ageing, thereby leading to the increased inflammation observed in the elderly^34^. In accordance, increased longevity has been reported in germ free mice^37^. Here we report both a decreased resident macrophage density and an increased neutrophil density with age. These changes during ageing in innate immune populations could be a contributing factor that may negatively affect tissue repair outcome in old mice, as evidenced by the attenuation of ageing hallmarks in liver that presented “youthful” resident macrophage density. Indeed, in patients that experienced a myocardial infarction or stroke, increased inflammatory parameters like Il-6 or TNF-RI were associated with older age^38^. Since resident macrophages have critical functions during tissue homeostasis and repair, in part through the swift removal of apoptotic cells and the secretion of cytokines and growth factors^39^, the reduction in resident macrophage numbers associated with an increased neutrophil infiltration could contribute to increased neutrophil activation within tissues, thereby leading to deleterious effects on tissue integrity^40^ and exacerbating age-induced tissue dysfunction.

## Methods

### Animals

*Csf1r*^*MeriCreMer*^ (ref ^41^), *Flt3*^*Cre*^ (ref ^42^), *Ticam*^*Lps2*^ (ref ^35^), *Rosa26*^*YFP*^ (ref ^43^) and *Rosa26*^*tdTomato*^ reporter (ref ^44^) mice have been previously described. *Csf1r*^*MeriCreMer*^ and *Csf1r*^*iCre*^ mice were on FVB background, other mice were on C57BL/6 (CD45.2) background. *Csf1r*^*MeriCreMer*^ mice were generated by J. W. Pollard. *Flt3*^*Cre*^ mice were generated by C. Bleul and provided by T. Boehm and S. E. Jacobsen. *Rosa26*^*tdTomato*^ (B6.Cg-Gt(ROSA)26Sortm9(CAG-tdTomato)Hze) and *Rosa2*6^YFP^ (B6.129X1-Gt(ROSA)26Sortm1(EYFP)Cos/J) were provided by S. Tajbakhsh and G. Eberl, respectively. Aged and Young C57BL/6JRj were purchased from Janvier labs. No randomization method was used and the investigators were blinded to the genotype of the animals during the experimental procedure. Results are displayed as mean ± s.e.m. All experiments included littermate controls and the minimum sample size used was 3. No statistical method was used to predetermine sample size. Animal procedures were performed in accordance with the Care and Use Committee of the Institut Pasteur (CETEA) guidelines and with their approval.

### Genotyping

PCR genotyping of *Ticam*^*Lps2*^ (ref ^35^), *Csf1r*^*MeriCreMer*^ and *Flt3*^*Cre*^ (ref ^20^) mice was performed according to protocols described previously.

### Fate-mapping of *Flt3*^*+*^ haematopoietic progenitors

For fate-mapping analysis of Flt3^+^ precursors, *Flt3*^*Cre*^ males (the transgene is located on the Y chromosome) were crossed to homozygous *Rosa26*^*YFP*^ reporter females. *Flt3*^*Cre*^ *Rosa*^*YFP*^ males were blood phenotyped. Animals with YFP labelling efficiency above 60% in the lymphocytes, monocytes and granulocytes were used for experiments and female littermates were used as Cre-negative controls.

### Pulse labelling of adult macrophages

For pulse labelling experiment of *Csf1r*^*+*^ macrophages in Young and Old mice, *Csf1r*^*MeriCreMer*^ *Rosa26*^*tomato*^ mice and Cre-negative littermates were injected intraperitoneally with 70 mg/kg of body weight tamoxifen (20mg/ml in corn oil, Sigma-Aldrich).

### Poly (I:C) acute and sustained inflammation experiments

For sustained inflammation experiments, mice were injected intraperitoneally with 5 mg/kg of saline or Poly (I:C) (Novus biologicals) every other day for one month. Mice were then sacrificed 48 hours, 2 weeks or 6 weeks after the last injection. For acute inflammation experiments, mice were injected intraperitoneally with 5 mg/kg of saline or Poly (I:C) received a total of 4 injections, two injections on alternate days, one week rest and two more injections on alternate days. Organs were harvested for flow cytometry and histology analyses 48 hours and 2 weeks after the last injection.

### Macrophage depletion using CSF1R inhibition

To deplete macrophages, mice were fed with the selective CSF1R inhibitor, PLX5622 (Plexxikon Inc.), for 3 days. Control and PLX5622 (300 ppm formulated in AIN-76A standard chow, Research Diets, Inc.) were purchased from Brogaarden APS.

### Processing of tissues for flow cytometry

Mice were killed by cervical dislocation and perfused by gentle intracardiac injection of 10 ml prewarmed (37°C) PBS 1X. Organs were weighted after harvesting. After being cut in small pieces, they were digested for 30 minutes at 37°C in PBS 1X containing 1mg/mL Collagenase D (Roche), 100 U/mL DNAse I (Sigma), 2.4 mg/mL Dispase (Invitrogen) and 3% FBS (Invitrogen). Epidermal sheets were separated from the dermis after incubation for 45 min at 37°C in 2.4 mg/ml of Dispase and 3% FBS and the epidermis was further digested for 30 min in PBS containing 1 mg/ml collagenase D, 100 U/ml DNase I, 2.4 mg/ml of Dispase and 3% FBS at 37°C. Tissues were then mechanically disrupted with a 2mL-syringe piston on top of 100 µm strainer to obtain a single cell suspension. For liver, hepatocyte depletion was performed by centrifugation of the whole liver cell suspension at 50g for 3 minutes. The supernatant was then collected for further staining of non-parenchymal liver cells. For liver, spleen and lung, red blood cell lysis was performed as previously described^45^.

### Flow cytometric analysis of adult tissues

Cells were centrifuged at 320g for 7 min, resuspended in 4°C FACS Buffer (PBS 1X, 2mM EDTA, 0.5% BSA), plated in multi-well round-bottom plates and immunolabelled for FACS analysis. After 15 min incubation with purified anti-CD16/32 (FcγRIII/II) diluted 1/50, antibody mixes were added and incubated for 30 min. When appropriate, cells were further incubated with streptavidin conjugates for 20 min. Samples were acquired using a Beckman Coulter Cytoflex S or a BD Fortessa cell analyser. All data was analysed using FlowJo 10.5.3 (BD). Cell density was calculated using the following formula: 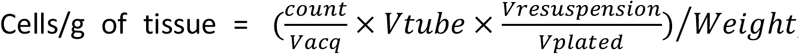, with Vacq: volume acquired; Vresuspension: volume added to the cell pellet after first centrifugation; Vplated: volume of cells plated; Vtube: total final volume; Weight: weight of the piece of tissue used for single-cell suspension preparation.

### EdU incorporation

EdU incorporation analysis was performed following the manufacturer’s instructions (BD EdU flow Kit). For steady state proliferation analysis in *Flt3*^*Cre*^ *Rosa*^*YFP*^ mice, 1 mg of EdU was injected intraperitoneally three times, every other day. In CSF1R inhibition (PLX) experiments, 1 mg of EdU was injected intraperitoneally 2 hours before sacrifice.

Liver single-cell suspensions were prepared as described above and stained with antibodies directed against CD45, CD11b, F4/80 and Tim4. Cells were then fixed, permeabilised and treated with DNAse according to manufacturer’s instructions and stained with the APC-EdU antibody and an anti-GFP antibody (Fischer Life).

**Table1:** Antibodies used for flow cytometry.

| Antibody | Clone | Conjugates |
| --- | --- | --- |
| CD45.2 | 104 | APC-Cy7 |
| CD11b | M1/70 | PE-Cy7 |
| F4-80 | BM8 | BV421 |
| TIM-4 | RMT4-54 | AF647or PE-Cy7 |
| CD64 | X54-5/7.1 FC | AF647 or PE-Cy7 |
| Ly-6C | HK1.4/AL-21 | APC, BV605 or FITC |
| Ly-6G | 1A8 | PE, or BV711 |
| Gr-1 | RB6-8C5 | PE or PerCP Cy5.5 |
| MHC-II (IA/IE) | M5/114.15.2 | AF647 |
| CD115 (Csf1r) | AF598 | AF647 or PE |
| CD3e | 145.2C11 | FITC or PE |
| CD19 | 1D3 | eF450 |

### RNA sequencing of Young and Old KCs

For RNA sequencing of Young (4 months) and Old (22-23 months) KCs, livers were processed as described before and KCs were then sorted with a BD FACS Aria III based on F4/80 and Tim4 expression. Labelling of cells was performed as described above using the following antibodies: CD45-APC-Cy7, CD11b-PE-Cy7, F4/80-BV421 and Tim4-APC. Gates were defined using unstained, single stained and fluorescence minus one (FMO) stained cells. 10 000 cells were directly sorted in 350 µL of RLT buffer (Qiagen), vigorously vortexed for disruption and spin down. RNA was subsequently extracted using the Qiagen RNeasy micro kit. Purified RNA was quantified and the RNA quality assessed using an Agilent Bioanalyzer (Agilent). Selected samples had an RNA Integrity Number > 6.2. Library preparation was performed using the ClonTech SMARTer Stranded RNA-SEq Kit V2 for mammalian pico RNA input (Takara) and RNA sequencing was performed using HiSeq 2500 sequencer with 50bp read lengths (Illumina).

#### Differential expression analysis

DESeq2 (v_1.22.2)^46^ Wald test was used to assess differential expression between groups. The input data consists of the raw count matrix where each row indicates the transcript, each column indicates the sample and each cell indicates the number of reads mapped to the transcript in the sample. After independent DESeq2 filtering^46^, P-values for selected genes were adjusted for multiple comparisons correction using Benjamini–Hochberg correction for FDR at threshold of P<0.05 and no minimum LFC threshold was applied.

#### Gene set enrichment analyses

Gene set enrichment analysis (GSEA) (functional scoring method (FSC)^47^) was conducted using the runGSA() function in piano R package^48^ was performed on the ranked list of genes. For this, the result table from DESeq2 was ordered by the –log10(adjusted p-value) from the Wald statistic test. Mouse Gene Symbol and Entrez IDs were added using the org.Mm.eg.db (v_3.7.0) and AnnotationDbi (v_1.44.0) Bioconductor Packages, using the Ensembl genes as keys, and duplicated entries identifiers were then collapsed by keeping the one with the lowest adjusted p-value. Gene set collections from the mouse version of the Molecular Signatures Database MSigDB v6 were downloaded from http://bioinf.wehi.edu.au/software/MSigDB/

#### Visual representation

Heatmap were made using R package pheatmap (v_1.0.12) and other visual representations (barplot and PCA biplot) were made using R package ggplot2 (v_2_3.1.0)^49^.

#### R Session info

All the analyses have been performed using R^50^ version 3.5.1 (2018-07-02), running under: OS X macOS Sierra 10.12.6 on x86_64-apple-darwin15.6.0 (64-bit) platform.

### Bead phagocytosis assay

Young (2-4 months) and Old (19 months) mice were injected intravenously twice with 2μm red fluorescent polymer microspheres (100µl of 0.2% solid beads in NaCl 0.9%) (FluoroMax, Molecular Probes). Livers were then analysed 24 hours after the last injection.

### Histology

Quantification of neutrophils and resident macrophages was performed in cryo-sections from Young (2-4mO) and Old (20-25mO) *Flt3*^*Cre*^ *Rosa*^*YFP*^ mice. Half of the organs were fixed overnight in 4% paraformaldehyde (Sigma) under agitation. After fixation, organs were washed in PBS 1X and incubated overnight in 30% sucrose, then embedded in OCT compound (Sakura Finetek). Cryoblocks were cut at a thickness of 12-14 µm and then blocked with PBS 1X containing 5% normal goat or donkey serum (Invitrogen); 0.2% BSA(w/v), 0.3% Triton X 100 for 1 hour at room temperature. Samples were incubated overnight at 4°C with primary antibodies (goat anti mouse MPO (1:100, Abcam); Armenian Hamster anti mouse CD31 (1:200, Abcam); rat anti Siglec-F (1:100, BD Pharmingen); rat anti mouse F4-80 (1:200, Bio-Rad); rabbit anti Ki67 (1:200, Abcam); rabbit anti GFP (1:300, Fischer Scientific); chicken anti GFP (1:300, Abcam) and rat anti PU.1 (1:200, R&D), rabbit polyclonal anti Desmin (1:100, abcam), goat polyclonal anti MCSF (1:50, bio-techne), sheep polyclonal anti IL-34 (1:20, bio-techne)) in PBS 1X; 1% serum; 0.2% BSA and 0.3% Triton X 100. After washing with PBS 1X-0.3% Triton X 100, slides were incubated for 2h at room temperature with DAPI (1:1000, Invitrogen) and appropriate secondary antibodies (goat anti Armenian Hamster AF647 and AF555 (1:500, Jackson Immuno); goat anti Rat AF555 and AF647, goat anti Rabbit AF488, goat anti Rabbit AF555, goat anti Chicken AF488 and goat anti Chicken AF555 (1:500, Invitrogen); chicken anti Goat AF647 (1:500, Thermo Fischer)) in PBS 1X; 0.2% BSA and 0.3% Triton X 100. Samples were then mounted with Prolong Gold antifade medium and imaged using an inverted Leica SP8 confocal microscope with 20 x/1.3 and 40x/1.3 (oil) objectives or a slide scanner (Olympus VS120) with 20x and 40x objectives.

Liver and lung neutrophils were identified as MPO positive cells and were quantified in nine fields of view (FoV, 0.3mm^2^) per animal (n=3 per age group), acquired with the 20X objective from a confocal microscope. The intra-tissular fraction was defined as MPO positive cells not colocalising within CD31+ sinusoids or large vessels. Liver macrophages were identified as F4-80+ cells and were quantified in six FoV (1mm^2^) per animal (n=3 per age group), acquired with the 20X objective from a slide scanner (Olympus VS120). Proliferating macrophages were identified by nuclear Ki67+ staining and YFP positivity of macrophages was also scored after staining with anti GFP antibody. Quantifications were performed blindly to age by two independent observers. For the niche component analysis, Stellate cells were identified as Desmin positive cells in liver sections from WT, *Flt3*^*Cre*^ *Rosa*^*YFP*^ and *Ticam*^*Lps2*^. Tunel staining was performed with the In Situ Cell Death detection kit (Sigma) as per the manufacturer’s instructions.

SA β-Gal staining on liver cryo-sections was performed as previously published^51^. Briefly, 12-20 µm liver sections from WT and TRIF Lps2 mice were stained as follows : sections were fixed 10 min in PBS 2% v/v PFA, 0,2% v/v Glutaraldehyde, washed three times in PBS and incubated 16 hours in a staining solution (40 mM Citric Acid at pH 6, 150 mM NaCl, 5 mM K3Fe(CN)6, 5 mM K4Fe(CN)6, 2 mM MgCl2 and 1mg/mL X-Gal in H2O). Sections were then washed 3×5 min in PBS and fixed in 4% formaldehyde. After washing 3×5 min in PBS, they were counterstained with fast-red for 10-15 seconds, before being rinsed under tap water. Sections where then incubated with Ethanol 95% for 5 min, Ethanol 100% for 2×5 min, and mounted in Eukitt mounting medium. Brightfield Images were then acquired with an Olympus VS120 Slide Scanner at a 20X magnification. Quantifications were done using the Fiji open-source software. Images were thresholded using a global thresholding algorithm (MaxEntropy) and the fraction of stained area was quantified. Eight to ten 0,34mm^2^ FOV were quantified per animal, in 5 animals per group. For the evaluation of lipid deposition, 12 µm liver sections from WT and TRIF Lps2 mice were stained with Oil Red O (Sigma) as per the manufacturer’s instructions. Brightfield Images were acquired with an Olympus VS120 Slide Scanner at a 40X magnification, within 24 hours after the staining. Quantifications were done using the Fiji open-source software. Images were thresholded using a local thresholding algorithm (Phansalkar with a 50 pixel radius) and the fraction of stained area was quantified. Eight to ten 0,34mm^2^ FOV were quantified per animal in 5 animals per group minimum.

### Statistical Analyses

Statistical analyses were performed using GraphPad Prism 7.0a (GraphPad software). Non-parametric t-test (Mann-Whitney) and non-parametric ANOVA (Kruskall Wallis) tests were performed, followed by a post-hoc Dunn comparison tests when significant differences were found.

## Data availability

All data generated or analysed during this study are included in this article (and its supplementary information files). Sequencing data sets described in this work will be submitted to the Gene Expression Omnibus (GEO) repository before final publication.

## Acknowledgments

We thank Ana Cumano, François Schweisguth, Geneviève Millon and Molly Ingersoll for reading the manuscript, Frédérique Vernel-Pauillac for the *Trif* mutant animals and the ageing and longevity Pasteur transversal program for scientific support. We thank the Plateforme de Cytometrie (Sophie Novault) for advice and help with cell sorting, and the Transcriptome and EpiGenome, BioMics, Center for Innovation and Technological Research of the Institut Pasteur for NGS. This work was supported by recurrent funding from the Institut Pasteur, the CNRS, and Revive (Investissement d’Avenir; ANR-10-LABX-73) and by an ERC investigator award (2016-StG-715320) from the European Research Council to E.G.P. E.G.P. also acknowledges financial support from the Fondation Schlumberger (FRM FSER 2017) and the Emergence(s) program from Ville de Paris (2016 DAE 190). K.A. was supported by a fellowship from the Complexité du Vivant doctoral school. LK and S.M. are supported by Revive.

## Author contributions

EGP proposed the concept, designed the experiments. EGP and KA wrote the paper. JSC and KA performed and analysed most experiments. DO, CK, PD and YL performed experiments. SM performed RNA sequencing analysis. CC and HL performed SA β-Gal experiments. LK provided aged cohorts and CW provided *Ticam*^*Lps2*^ young and aged mice. All authors read and agreed on the manuscript.

## Author information

The authors declare no competing interests. Correspondence and requests for materials should be addressed to.

## Notes

### Competing Interest Statement

The authors have declared no competing interest.

