## supplemental figures for "Inflammation drives age-induced loss of tissue resident macrophages"

### Sup Figures Ade, Saenz Coronilla et al.

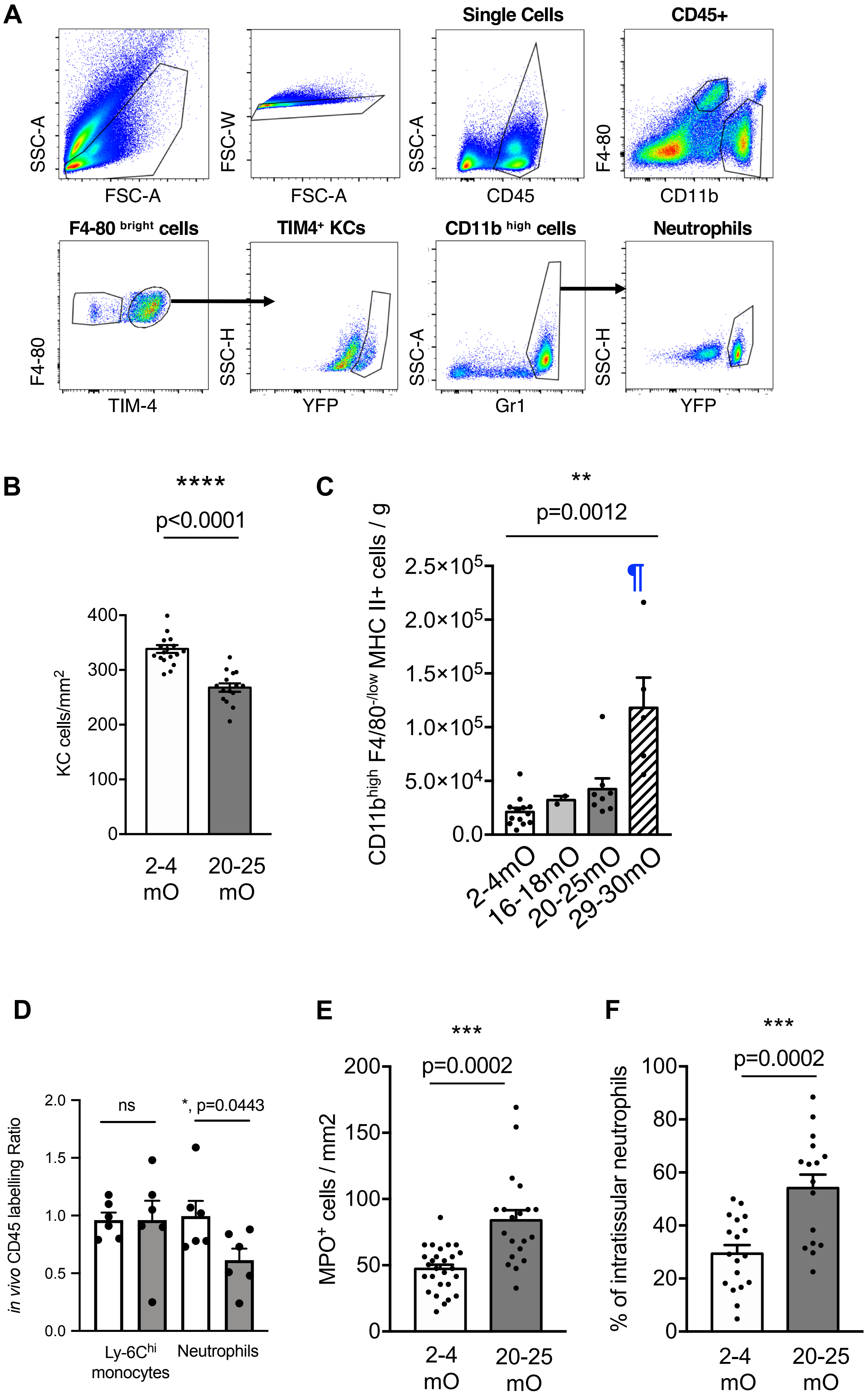

A
SSC-A
FSC-A
FSC-W
FSC-A
Single Cells
SSC-A
CD45
CD45+
F4-80
CD11b
F4-80 bright cells
F4-80
TIM-4
TIM4+ KCs
SSC-H
YFP
CD11b high cells
SSC-A
Gr1
Neutrophils
SSC-H
YFP
B
****
p<0.0001
KC cells/mm2
400
300
200
100
0
2-4 mO
20-25 mO
C
**
p=0.0012
CD11b high F4/80 -/low MHC II+ cells / g
2.5×10^5
2.0×10^5
1.5×10^5
1.0×10^5
5.0×10^4
0.0
¶
2-4mO
16-18mO
20-25mO
29-30mO
D
in vivo CD45 labelling Ratio
2.0
1.5
1.0
0.5
0.0
ns
*, p=0.0443
Ly-6Chi monocytes
Neutrophils
E
***
p=0.0002
MPO+ cells / mm2
200
150
100
50
0
2-4 mO
20-25 mO
F
***
p=0.0002
% of intratissular neutrophils
100
80
60
40
20
0
2-4 mO
20-25 mO

**Fig. S1: Age-associated changes in liver innate immune cells.**

**A**, Gating strategy for liver Kupffer cells (KC) and neutrophils. **B**, Number of F4/80$^+$ liver KC per mm$^2$ in cryo-sections from Young (2-4 mO) and Old (20-25 mO) mice. Mean ± s.e.m; Mann-Whitney test; n=3 per age group. **C**, Number of CD11b$^{high}$ F4/80$^{-/low}$ MHC-II$^+$ cells per g of liver from Young (2-4mO; n=13) and Old 16-18mO (n=3), 20-25mO (n=8) and 29-30mO (n=5) animals. Mean ± s.e.m; one-way ANOVA (Kruskal Wallis) and post-test Dunn multiple comparison test against Young group: ¶, p<0.001. **D**, ratio of CD45 labelling in the tissue compared to the labelling of the circulating counterparts, after in vivo injection of CD45 antibody 2 minutes before sacrifice. **E**, Number of MPO$^+$ neutrophils per mm$^2$ in liver and **F,** Frequency of intra-tissular liver neutrophils in cryo-sections from Young (2-4 mO) and Old (20-25 mO) mice. Mean ± s.e.m; Mann-Whitney test; n=4 per age group.

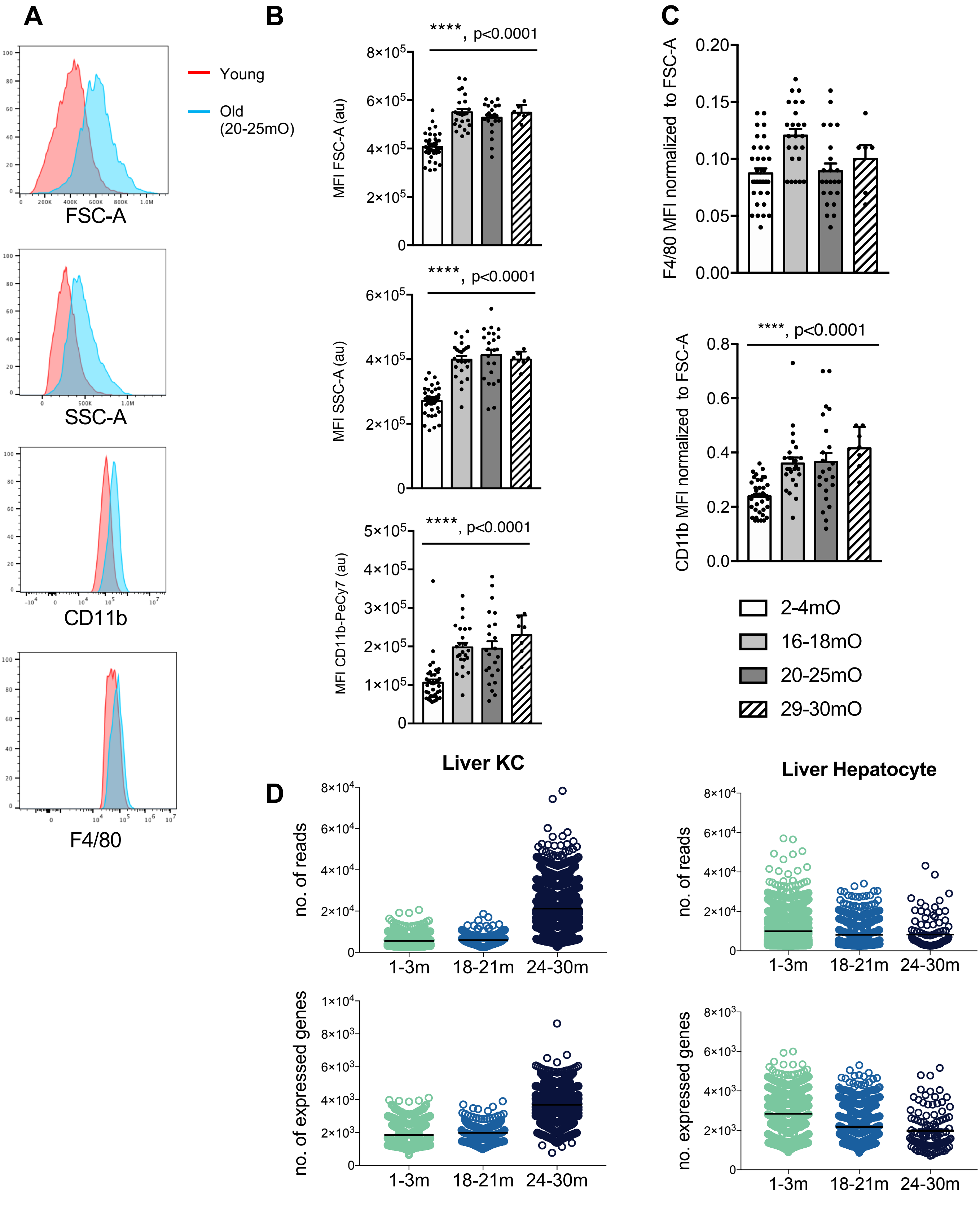


**Fig. S2: Age-associated morphological changes in liver resident macrophages.**

**A**, Representative histograms of SSCA (granularity), FSC-A (size), CD11b and F4/80 expression in Young (red line) and Old (20-25 mO; blue line) liver Kupffer cells (KC). **B**, Median Fluorescent Intensity (MFI) of SSC-A and FSC-A and **C**, of F4-80 and CD11b expression normalized to size in Young (2-4 mO, n=40) and Old 16-18mO (n=24), 20-25mO (n=23) and 29-30mO (n=7) KCs. Mean ± s.e.m; one-way ANOVA (Kruskal Wallis). **D**. Number of reads for the KC and Hepatocyte clusters, extracted from the Tabula Muris Senis consortium data set.

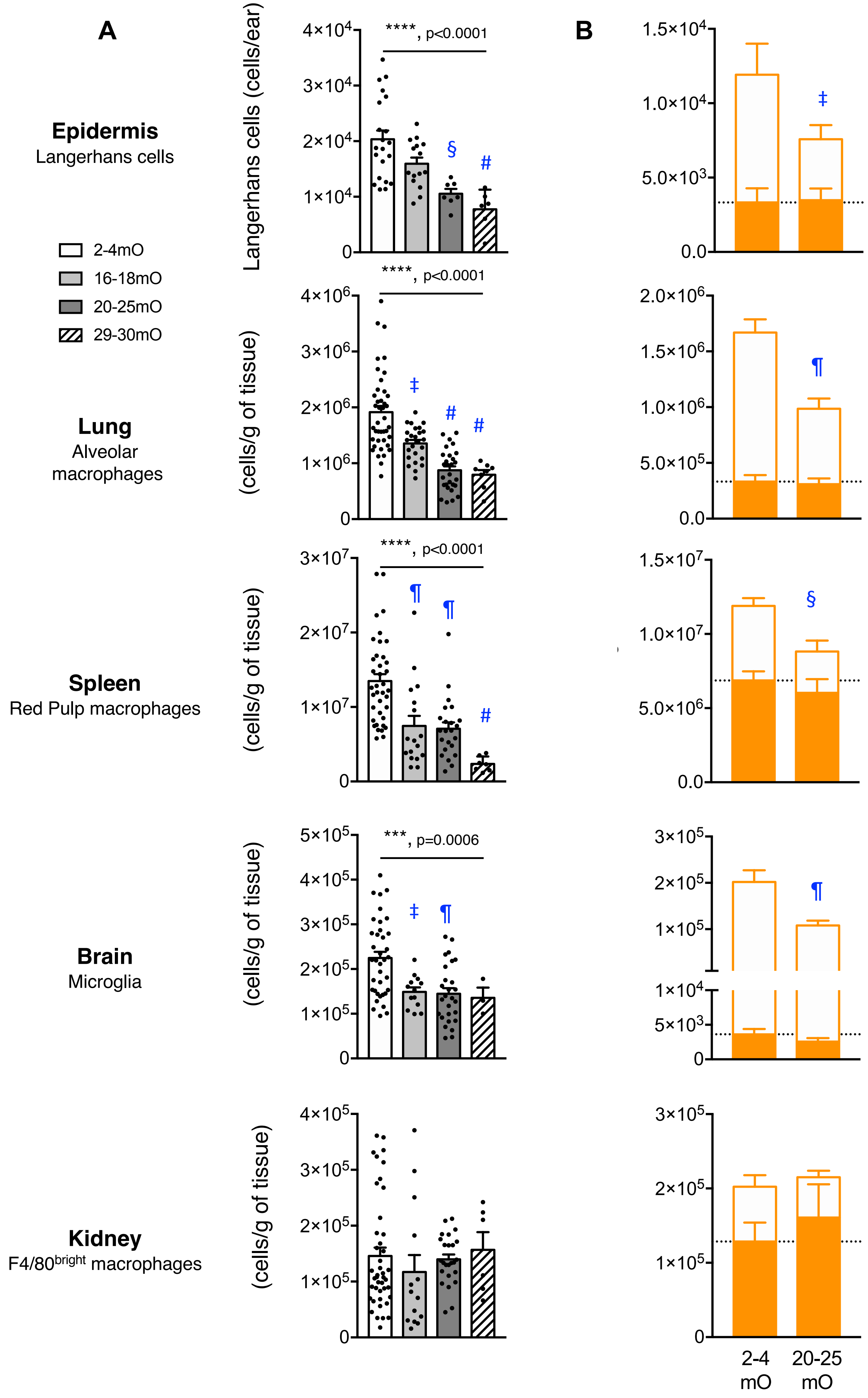
A
B
Epidermis
Langerhans cells
Lung
Alveolar macrophages
Spleen
Red Pulp macrophages
Brain
Microglia
Kidney
F4/80bright macrophages
2-4mO
16-18mO
20-25mO
29-30mO
Langerhans cells (cells/ear)
(cells/g of tissue)
****, p<0.0001
***, p=0.0006
§
#
‡
¶
2-4 mO
20-25 mO

**Fig. S3: Loss of HSC-independent resident macrophages with ageing in most tissues except in the kidney.**

**A**, Number of resident macrophages per ear for epidermis (Langerhans cells) and per g of tissue for lung (Siglec-F$^{+}$ F4/80$^{bright}$ CD64$^{+}$), brain (F4/80 $^{+}$), spleen (F4/80$^{bright}$ CD64$^{+}$) and kidney (F4/80$^{bright}$ CD64$^{+}$) from Young (2-4mO; n=19-41) and Old 16-18mO (n=4-24), 20-25mO (n=7-24) and 29-30mO (n=3-11) animals. Mean ±s.e.m; one-way ANOVA (Kruskal Wallis) per organ and Dunn multiple comparison test against Young group: #, $p<0.0001$; ¶, $p<0.001$; §, $p<0.01$; ‡, $p<0.05$. **B**, Number of (*Flt3*$^{Cre}$)-YFP$^{+}$ and (*Flt3*$^{Cre}$)-YFP$^{neg}$ resident macrophages per ear for epidermis and per g of tissue for lung, brain spleen and kidney. Dashed lines represent the mean density of (*Flt3*$^{Cre}$)-YFP$^{+}$ resident macrophages; mean ±s.e.m; Young, n=9 and Old, n=8; Mann Whitney test of Young versus Old YFP$^{neg}$ cells: ¶, $p<0.001$; §, $p<0.01$; ‡, $p<0.05$.

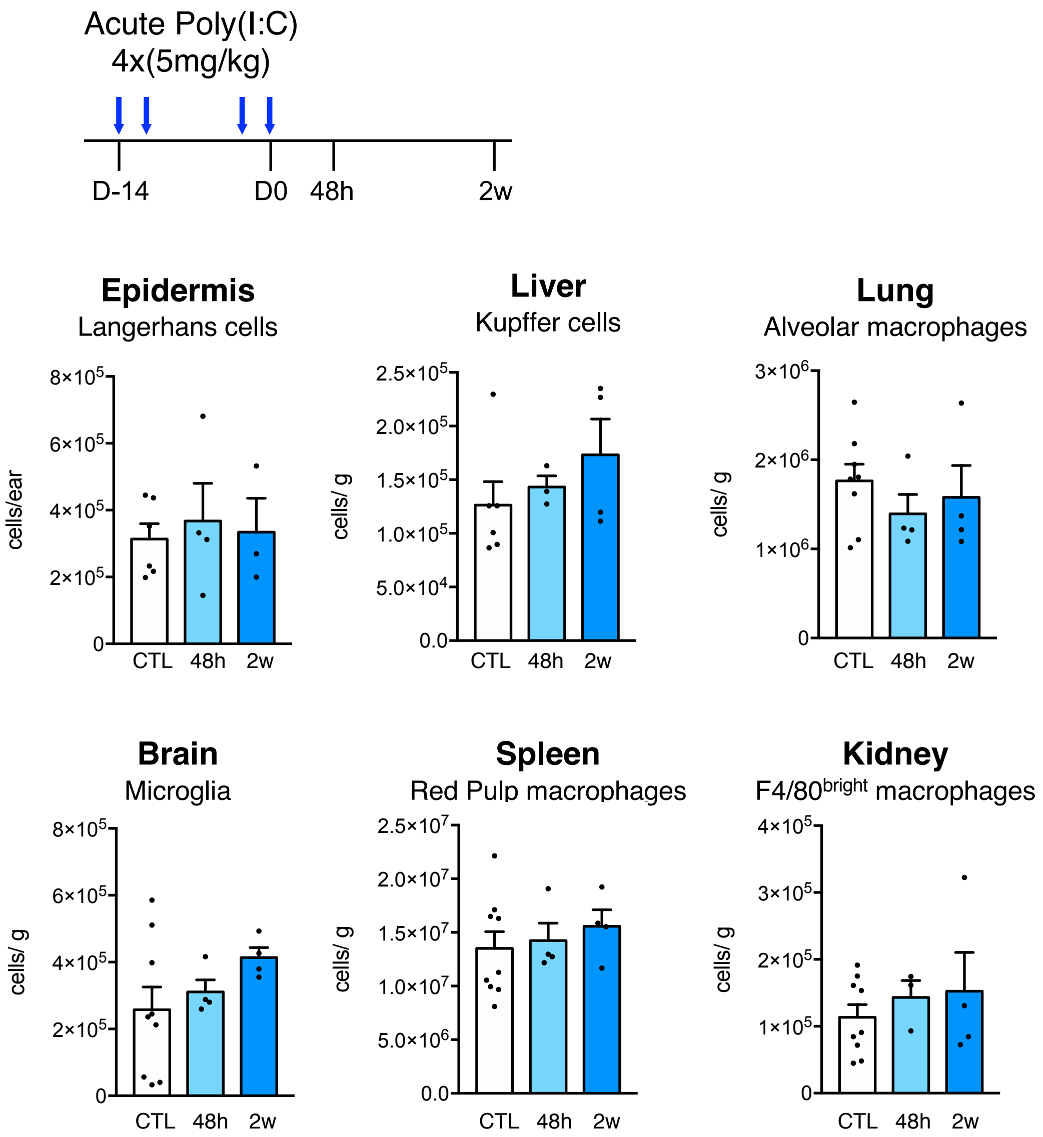


**Fig. S4: Macrophage and Neutrophil density after acute Poly(I:C)-induced inflammation, respectively.**

Analysis 48h and 2 weeks (2w) after acute (4 injections within ten days) saline (control, CTL) or Poly(I:C) injections. Number of resident macrophages per ear (Langerhans cells) and per g of liver (Kupffer cells), lung (alveolar macrophages), brain (microglia), spleen (red pulp macrophages) and kidney. Mean ±_s.e.m; saline control, n=6; 48h, n=4 and 2w, n=4.

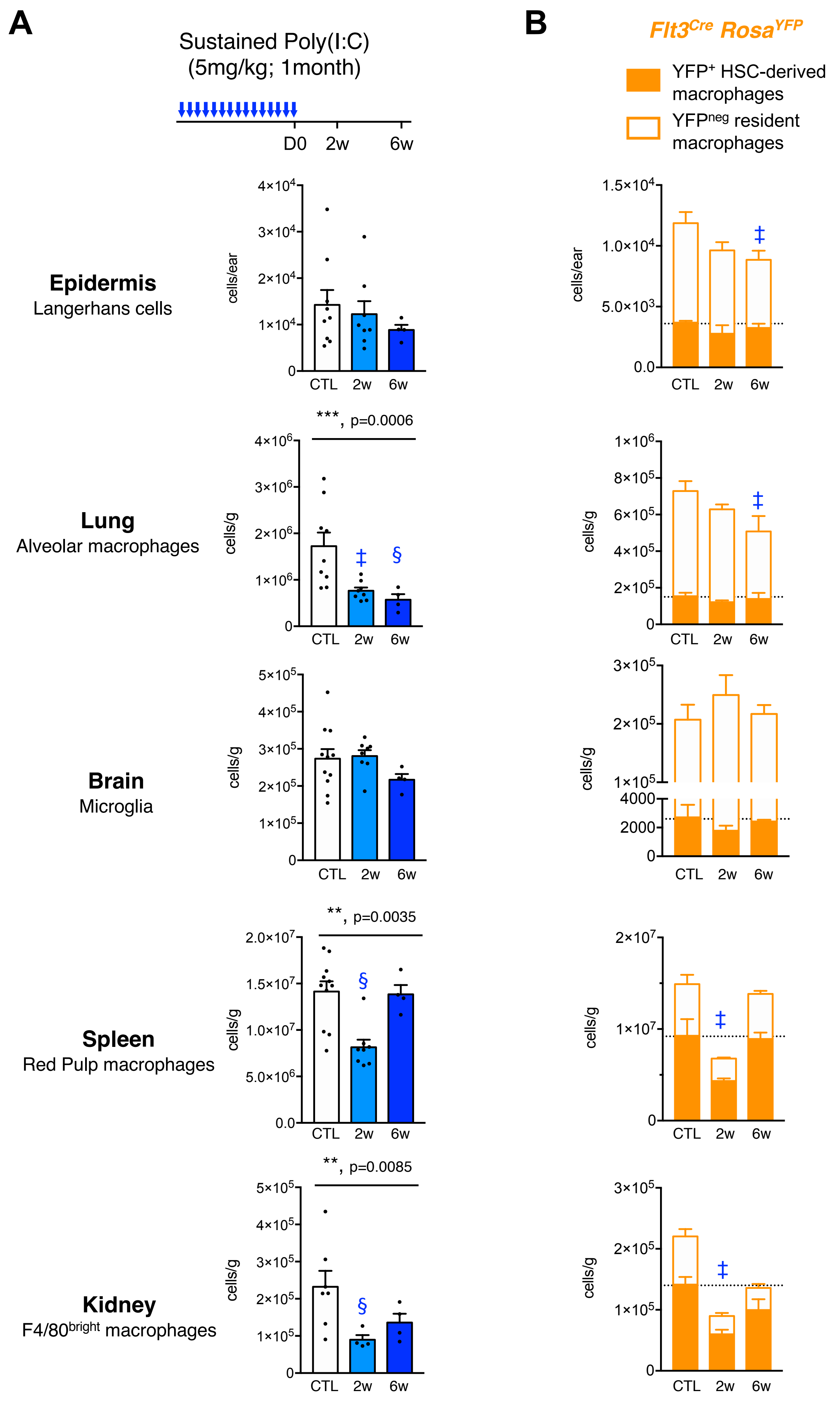
A
Sustained Poly(I:C)
(5mg/kg; 1month)
D0
2w
6w
B
Flt3Cre RosaYFP
YFP+ HSC-derived macrophages
YFPneg resident macrophages
Epidermis
Langerhans cells
Lung
Alveolar macrophages
Brain
Microglia
Spleen
Red Pulp macrophages
Kidney
F4/80bright macrophages
cells/ear
cells/g
***, p=0.0006
**, p=0.0035
**, p=0.0085
CTL
2w
6w

**Fig. S5 : Resident macrophage density after repeated poly(I:C) inflammation in epidermis, lung, brain, spleen and kidney**

**A**, Analysis 2 weeks (2w) and 6 weeks (6w) after sustained challenge with every other day injections for a month of saline (control, CTL) or Poly(I:C) in Young adult mice. Number of resident macrophages per ear for epidermis and per g of tissue for lung, brain spleen and kidney. Mean ±_s.e.m; one-way ANOVA (Kruskal Wallis) per organ and post-test Dunn multiple comparison test against saline group: §, $p<0.01$; ‡, $p<0.05$; saline control, n=9-11; sustained Poly(I:C) 2w, n=4-8 and 6w, n=4. **B**, Analysis 2w and 6w after sustained saline (control, CTL) or Poly(I:C) injections in Young $Flt3^{Cre}$ $Rosa^{YFP}$ mice. Number of ($Flt3^{Cre}$)-$YFP^{+}$ and ($Flt3^{Cre}$)-$YFP^{neg}$ resident macrophages per ear for epidermis and per g of tissue for lung, brain spleen and kidney. Dashed lines represent the mean density of ($Flt3^{Cre}$)-$YFP^{+}$ resident macrophages in saline controls, mean ±_s.e.m; one-way ANOVA (Kruskal Wallis) per organ and Dunn multiple comparison test against $YFP^{neg}$ cells in saline group: ‡, $p<0.05$; saline control, n=3; chronic Poly(I:C) 2w, n=4 and 6w, n=4.

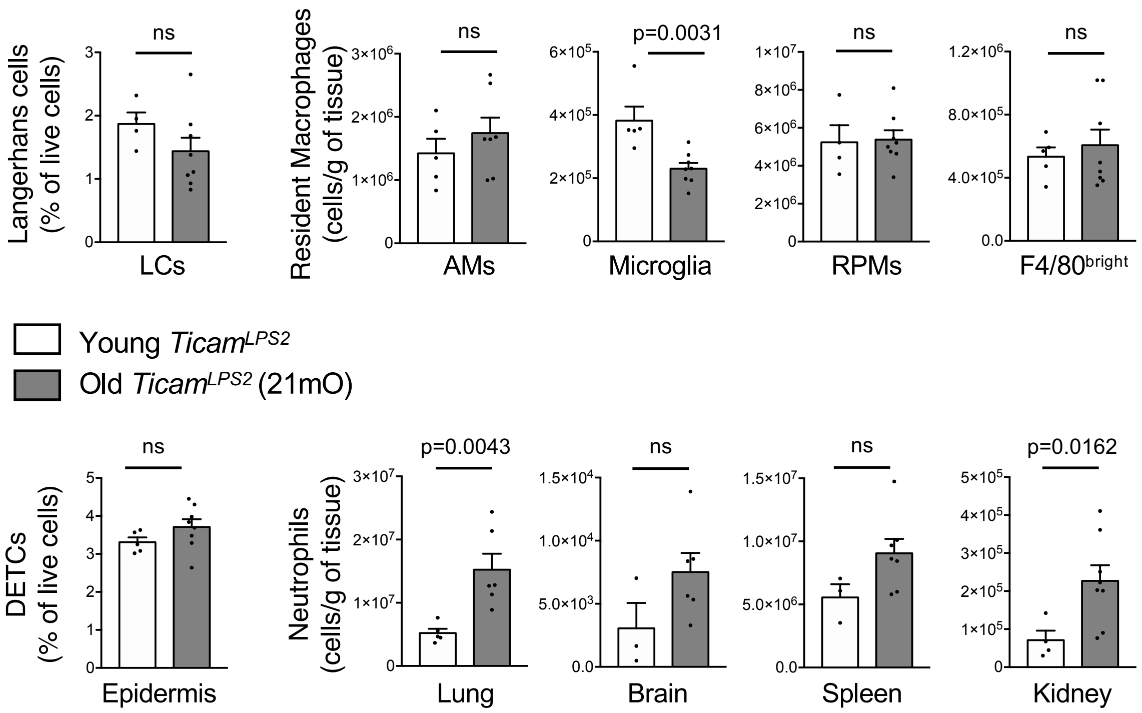


**Fig. S6: Neutrophil and macrophage density in epidermis, lung, brain, spleen and kidney of *Ticam$^{Lps2}$* mutants**

**A**, Frequency of neutrophils (orange), Gr1$^{-}$ and Gr1$^{+}$ monocytes (green and red, respectively) and lymphocytes (purple) in blood from Young (5 mO, n=5) and Old (21mO, n=8) *Ticam$^{Lps2}$* mutant animals. **B**, Frequency of epidermal Langerhans cells (LCs) and number of resident macrophages per g of lung (alveolar macrophages, AMs), brain microglia, spleen (red pulp macrophages, RPMs) and kidney (F4/80$^{bright}$ CD64$^{+}$ macrophages) and **C,** Frequency of dendritic epidermal T cells (DETCs) and number of neutrophils per g of lung, brain spleen and kidney from Young (5 mO, n=5) and Old (21mO, n=8) *Ticam$^{Lps2}$* mutant animals. Mean ± s.e.m; Mann-Whitney test.
